## Supplemental file 1 for "Pulsed ultrasound promotes secretion of anti-inflammatory extracellular vesicles from skeletal myotubes via elevation of intracellular calcium level": Supplemental material 1_Yamaguchi.docx

**Supplemental material 1-1**

List of specific miRNAs in the control group

miRNA

miR-7043-3p

miR-3087-3p

miR-8112

miR-485-3p

miR-185-3p

miR-7648-3p

let-7i-3p

miR-1960

miR-3061-3p

miR-3061-5p

miR-494-3p

miR-1934-5p

miR-1941-5p

miR-6914-3p

miR-6956-3p

miR-431-3p

miR-205-5p

miR-187-5p

miR-6948-3p

miR-6966-3p

miR-3473h-5p

miR-323-3p

miR-8103

miR-328-5p

miR-382-5p

miR-504-5p

miR-8105

miR-23a-5p

miR-6928-3p

**Supplemental material 1-2**

List of specific miRNAs in the ultrasound group

miRNA

miR-33-5p

miR-449a-5p

miR-467e-5p

miR-1929-5p

miR-1963

miR-1966-5p

miR-25-5p

miR-196b-3p

miR-700-5p

miR-8118

miR-12193-5p

miR-130b-3p

miR-7660-3p

miR-33-3p

miR-146b-3p

miR-675-5p

miR-3095-5p

let-7g-3p

miR-138-5p

miR-450b-5p

miR-324-3p

miR-743b-5p

miR-669a-5p

miR-669a-5p

miR-669a-5p

miR-669a-5p

miR-669a-5p

miR-669a-5p

miR-669a-5p

miR-669a-5p

miR-669a-5p

miR-669a-5p

miR-669a-5p

miR-669a-5p

miR-669p-5p

miR-669p-5p

miR-130a-5p

miR-6994-3p

miR-124-5p

let-7j

miR-542-5p

miR-7653-5p

miR-6901-5p

miR-452-3p

miR-7065-3p

miR-147-5p

miR-212-3p

miR-138-2-3p

miR-31-3p

miR-129-1-3p

miR-1930-5p

miR-3082-3p

miR-345-5p

miR-10a-3p

miR-744-3p

miR-1306-5p

miR-6945-3p

miR-153-5p

miR-324-5p

miR-298-3p

miR-7083-5p

miR-181a-1-3p

miR-7685-3p

miR-135b-5p

miR-150-5p

miR-1955-5p

miR-485-5p

miR-6948-5p

miR-465a-5p

miR-301a-5p

miR-344c-3p

miR-7021-5p

miR-223-5p

miR-3094-3p

miR-488-3p

miR-370-5p

miR-677-5p

miR-5132-5p

miR-28c

miR-200a-5p

miR-3060-3p
