## Supplemental file 2 for "Pulsed ultrasound promotes secretion of anti-inflammatory extracellular vesicles from skeletal myotubes via elevation of intracellular calcium level": Supplemental material 2_Yamaguchi.pptx

### Slide 1
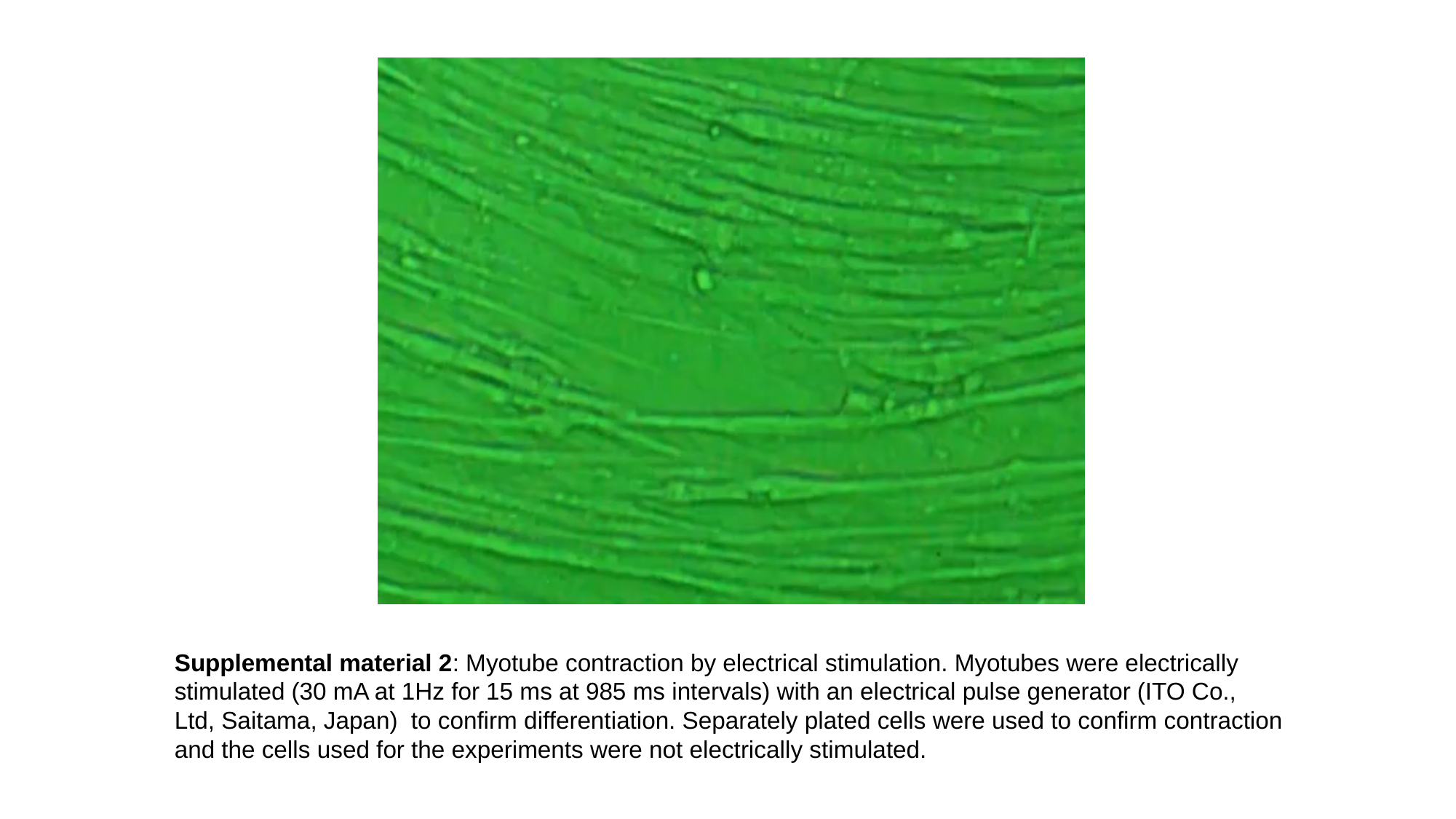

Supplemental material 2: Myotube contraction by electrical stimulation. Myotubes were electrically
stimulated (30 mA at 1Hz for 15 ms at 985 ms intervals) with an electrical pulse generator (ITO Co.,
Ltd, Saitama, Japan) to confirm differentiation. Separately plated cells were used to confirm contraction
and the cells used for the experiments were not electrically stimulated.
