## Supplemental file 3 for "Pulsed ultrasound promotes secretion of anti-inflammatory extracellular vesicles from skeletal myotubes via elevation of intracellular calcium level": Supplemental material 3_Yamaguchi.docx

Sequences for qPCR primers.

| Primer | Sequences |
| --- | --- |
| *Gapdh*-Forward | 5'-CCAATGTGTCCGTCGTGGATCT-3' |
| *Gapdh*-Reverse | 5'-GTTGAAGTCGCAGGAGACAACC-3' |
| *Il-1b*-Forward | 5'-GCCTTGGGCCTCAAAGGAAAGAA-3' |
| *Il-1b*-Reverse | 5'-ATTGCTTGGGATCCACACTCTCC-3' |
| *Il-6*-Forward | 5'-ACAAAGCCAGAGTCCTTCAGAGA-3' |
| *Il-6*-Reverse | 5'-TTGGATGGTCTTGGTCCTTAGCC-3' |
